# Beyond oscillations - A novel feature space for characterizing brain states

**DOI:** 10.1101/2024.04.17.589917

**Authors:** Elio Balestrieri, Nikos Chalas, Christina Stier, Jana Fehring, Cristina Gil Ávila, Udo Dannlowski, Markus Ploner, Joachim Gross

## Abstract

Our moment-to-moment conscious experience is paced by transitions between states, each one corresponding to a change in the electromagnetic brain activity. One consolidated analytical choice is to characterize these changes in the frequency domain, such that the transition from one state to the other corresponds to a difference in the strength of oscillatory power, often in pre-defined, theory-driven frequency bands of interest. Today, the huge leap in available computational power allows us to explore new ways to characterize electromagnetic brain activity and its changes.

Here we leveraged an innovative set of features on an MEG dataset with 29 human participants, to test how these features described some of those state transitions known to elicit prominent changes in the frequency spectrum, such as eyes-closed vs eyes-open resting-state or the occurrence of visual stimulation. We then compared the informativeness of multiple sets of features by submitting them to a multivariate classifier (SVM).

We found that the new features outperformed traditional ones in generalizing states classification across participants. Moreover, some of these new features yielded systematically better decoding accuracy than the power in canonical frequency bands that has been often considered a landmark in defining these state changes. Critically, we replicated these findings, after pre-registration, in an independent EEG dataset (N=210).

In conclusion, the present work highlights the importance of a full characterization of the state changes in the electromagnetic brain activity, which takes into account also other dimensions of the signal on top of its description in theory-driven frequency bands of interest.

## Introduction

Our everyday experience unfolds in an endless sequence of states: attentive, hungry, drowsy, startled… Each of these states coincides with characteristic behavioral outcomes as well as with a complex pattern of electrophysiological brain activity.

Research has long focused on the spectral features of these internal “brain states’’. Seminal EEG works showed that the transitions from rest to being engaged in a task (1), or from wakefulness to sleep or anesthesia (2,3) are well described by prominent changes in canonical frequency bands, such as alpha (8-13 Hz) and delta (1-4 Hz). Since then, brain oscillations have proven to be ubiquitous in characterizing diverse brain states (4), showing that shifts in oscillatory power pace movement planning and execution (5,6), precede perceptual decision-making outcomes (7–9), and play a critical role in the maintenance of items in working memory (10,11). Brain states thus sit at the nexus between physiology and behavior (12), and understanding their widely distributed pattern of activity is a goal both ambitious and crucial.

One way to tackle the complexity of brain states is to focus on single domains and cortical areas. A well-studied domain is the human visual system, where neural activity is shaped by the complex interplay of thalamo-cortical and cortico-cortical rhythms (13,14) pacing the transition from resting-states to active ones (15). Rhythmic activity is also posited to orchestrate both the flow of bottom-up information as well as the top-down modulation (16– 19). This is not the only perspective: recent accounts put to the test the importance of oscillations in visual information processing (20), and highlight the importance of other phenomena, such as aperiodic transients, in the feedforward flow of information (21).

Such a broadening of perspective is also methodologically grounded. In fact, various studies highlighted the limitations of exclusively concentrating on spectral parameters in time series analysis, particularly within specific frequency bands. The estimation of changes in oscillatory power is liable to confounders, including changes in aperiodic components (22) and the non-sinusoidal nature of oscillatory phenomena (23). Moreover, brain oscillations show clear signs of non-stationarity along a typical recording (24), or appear as isolated bursts whose interpretation could be distorted when averaging trials (25), or display a non-zero mean inducing baseline shifts (26). All these issues highlight the importance of expanding our analytic repertoire for the study of brain states beyond the traditional approaches based on oscillations in canonical frequency bands.

The aim of the present study is thus to present and validate novel ways to characterize brain states. We chose a set of brain states known for their emblematic and robust oscillatory patterns in the human visual cortex: absence of visual stimulation (resting state with eyes closed), resting state with eyes open, and task-related activity using a visual stimulation known to elicit strong gamma oscillations (27). We capitalized on previous works (28,29,30) to compose a novel feature space defining brain states along dimensions such as signal variability, complexity, aperiodic components, intrinsic timescales and entropy. We inferred the informativeness of these features in discriminating brain states by evaluating their classification accuracy. We further compared the classification accuracy of this aggregated set of features against the benchmark defined by power in canonical frequency bands, and we find a consistent advantage of the novel features. This result, corroborated by a pre-registered replication on an independent EEG dataset, encourages the exploration of new ways to characterize brain states.

## Results

We used a Support Vector Machine (SVM) to evaluate the brain states classification accuracy of 3 sets of features. The novel set of features, from hereafter “TimeFeats”, contained 41 descriptors of time series. Briefly, these included the first 4 moments of probability distributions, measures of entropy (31,32), a set of wide purpose features called *catch22 (28)* and features previously used in M-EEG, such as aperiodic spectral components, Hurst exponent, Hjort parameters and zero crossings (30). The second set, from hereafter “FreqBands”, contained 6 features, defined as power estimates averaged in frequency bands: delta (1-4 Hz), theta (4-8 Hz), alpha (8-13 Hz), beta (13-30 Hz), low gamma (30-45 Hz) and high gamma (55-100 Hz). The third, named “fullFFT”, contained 100 features as power in every frequency from 1 to 100 Hz, with a frequency resolution of 1 Hz. The four states taken into consideration were resting state with eyes closed (EC), resting state with eyes open (EO), visual stimulation consisting of a sine wave grating (VS) and the prestimulus baseline before the stimulation (BSL). Since all of these states are known to elicit changes in visual cortices, we constrained our analysis to the cortical parcels classified as visual in the MEG dataset, and to the posterior channels in the EEG dataset.

### Benchmark of novel features’ accuracy

In the MEG dataset of 29 participants (see *Methods, Main Experiment data collection*) we evaluated the informativeness of the novel features (*TimeFeats*) by comparing their classification accuracy with the benchmarks offered either by the power in canonical frequency bands (*FreqBands*) or by the full spectral power (*fullFFT*), all of them based on source-localized MEG data. For both the classification between eyes closed vs eyes open (EC/EO) and baseline vs visual stimulation (BSL/VS), we adopted different partition schemes. In the *“within”* scheme, cross-validation is performed within each subject, yielding one classification accuracy value for each subject averaged for all the cross-validation folds. In the “*between”* scheme, the data for all subjects minus one is concatenated together and the test accuracy is computed on the left-out subject. This procedure is repeated for all subjects, leading one accuracy value for each. In the “*across*” scheme, 85% of data from each subject is concatenated in the training set, and 15% in the test set. A single accuracy value, exclusively for the test set is hence reported. Results are shown in figure 1.

**Figure 1.**
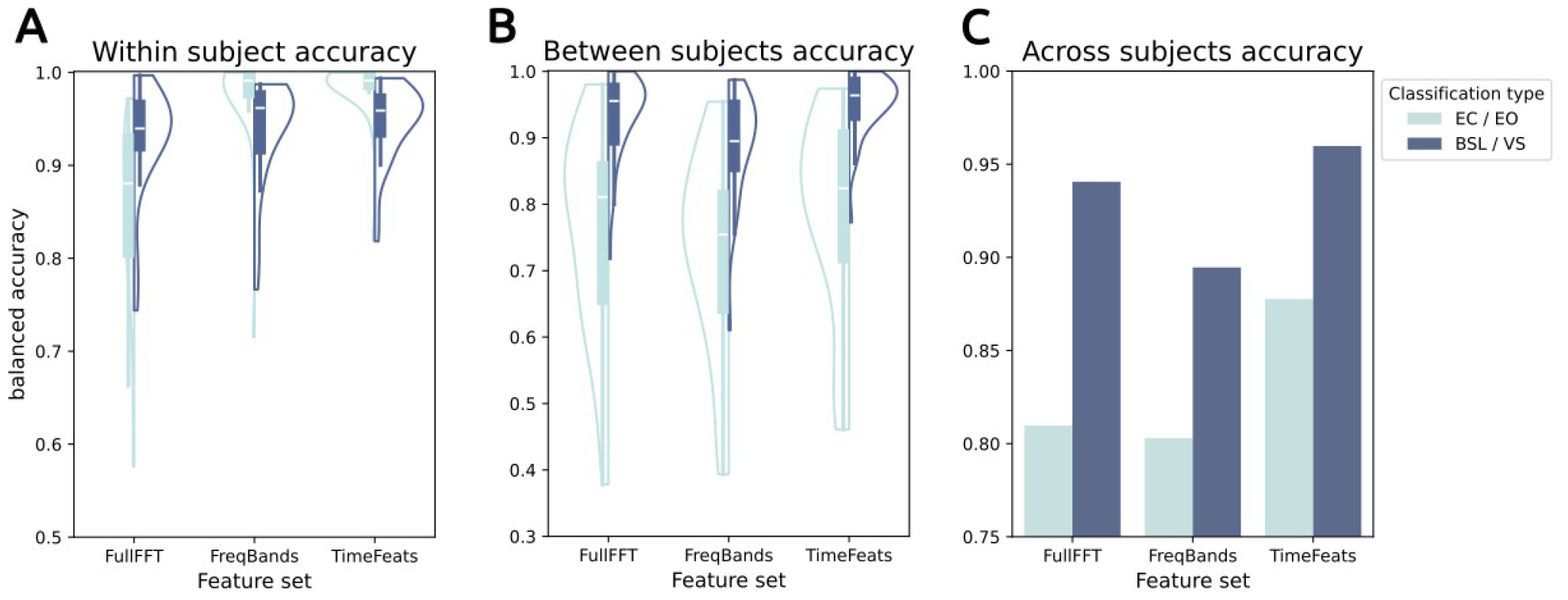
**A**: Within subject classification accuracy, pooled for all subjects. **B**: Between subjects classification accuracy. All subjects left out in every cross-validation fold are pooled together. **C**: Across subjects classification accuracy. Since all subjects are pooled in one train/test set, the analysis yields only one accuracy estimate per Feature set/Classification type.

In all conditions, we obtained a classification accuracy strongly above chance, and often above .9. The highest accuracy overall was achieved in the “within subject” scheme (figure 1A), where TimeFeats scored on average .982 in EC/EO and .946 in BSL/VS. A repeated measures ANOVA showed a significant effect of feature set (F_(2, 56)_=63.759, p<.001, η^2^=.227) and a significant interaction between feature set and classification type (F_(2, 56)_=43.908, p<.001, η^2^=.158). A repeated measure (rm) ANOVA comparing only the two top feature sets (FreqBands and TimeFeats) confirmed the feature set effect, hence favoring TimeFeats (F_(2, 28)_=6.217, p=.019, η^2^=.008). When evaluated separately for EC/EO and BSL/VS the difference between FreqBands and TimeFeats only approaches significance (EC/EO: t_(28)_=-2.049, p=.05, d=.256; BLS/VS: t_(28)_=-2.003, p=.055, d=.195), likely because of the ceiling effect in accuracy in this partition scheme.

The “between subject” scheme (figure 1B) yielded expectedly lower accuracy, although still strongly higher than chance, offering a proof of concept that all the features examined here generalize also to different subjects. Also here the top accuracy is reached from TimeFeats (EC/EO=.783; BSL/VS=.948) and a repeated measures ANOVA disclosed a significant effect of feature set via (F_(2, 56)_=54.535, p<.001, η^2^=.04) as well as an effect of the classification type (F_(2, 28)_=31.715, p<.001, η^2^=.332). This last result is of interest, suggesting that overall the features generalize better between participants during a task, rather than during resting state.

The second best feature set in this case is fullFFT (EC/EO=.756; BSL/VS=.931). This set is likely favored over FreqBands due to the higher number of features, which the classifier can leverage better given the great number of observations coming from concatenating multiple subjects together, an advantage absent in the “within subject” classification scheme. The pairwise comparison of accuracy between fullFFT and TimeFeats still favored the latter (EC/EO: t_(28)_=-3.187, p=.004, d=.292; BLS/VS: t_(28)_=-2.912, p=.007, d=.171).

The last partition scheme, “across subjects” does not allow inferential statistics, since all the subjects are pooled together in one single training/test datasets. Nonetheless, it confirms TimeFeats as the best performing set of features (EC/EO=.878; BSL/VS=.96).

### Comparison of single features

Our analysis compared features grouped in sets, showing that the novel features computed in the time domain perform better than the ones based on spectral power in canonical frequency bands. Nonetheless each of these sets pools together a great variety of features, whose specific contribution in the discrimination between brain states is yet to be unveiled. For this reason we further evaluated how every single feature, in every frequency band, is informative for discriminating between all the four states (EC/EO/BSL/VS) together (figure 2). This was achieved by computing classification accuracy for the 4 different states through a multiclass SVM. To reduce computation time, classification accuracy was computed uniquely “across subjects”.

**Figure 2.**
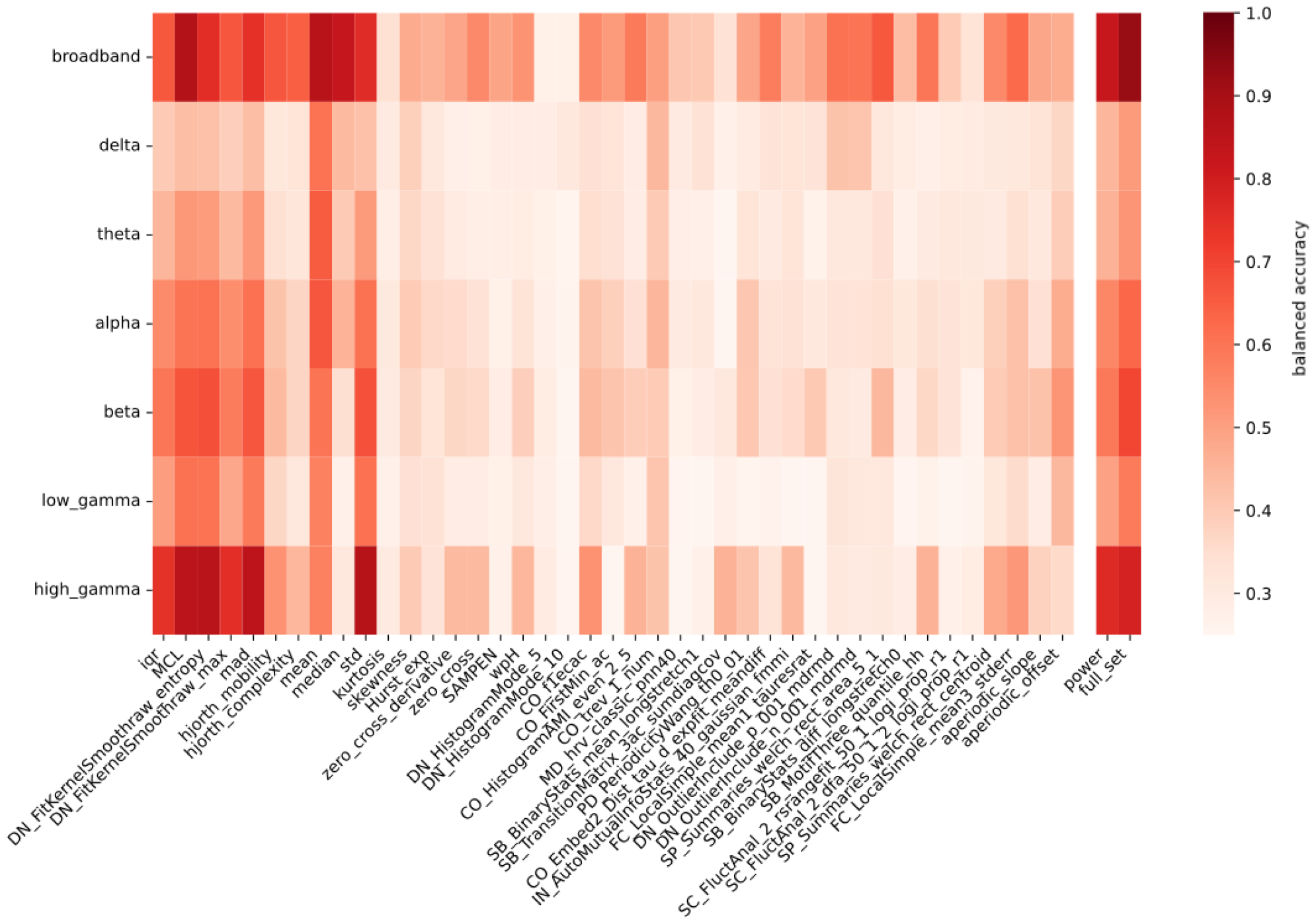
Across subjects classification accuracy for each feature, both computed on the broadband signal and separately in each frequency band. The first 41 columns correspond to single features from the TimeFeats set. The column “power” includes power in each frequency band, and in all these frequency bands combined (broadband). The last column (“full_set”) includes all the 41 time feats combined, either computed on the band-passed signal within frequency bands of interest or over the whole signal (broadband). Accuracy refers to the four classes classification: EC/EO/BSL/VS.

Once again, the full set of “TimeFeats” resulted in the best classification accuracy overall (figure 2: “full set” & “broadband” = .924). Interestingly, the second best performer was a single feature named Mean Curve Length (MCL), that with a classification accuracy of .873 also exceeded the power in all frequency bands (.826).

The most informative frequency band for the classification is high gamma, a result to be expected given the states considered: both the passage from eyes closed and eyes open, as well as the visual stimulation used in this experiment are characterized by an increase in broadband gamma power. What is intriguing is that, within the high gamma band, the classification accuracy scored by power (.764) was substantially lower than the one of several other features, including standard deviation (std) (.863), (MCL=.851), DN_FitKernelSmoothraw_entropy (.86) and mean absolute deviation (mad) (.85).

We further aimed to seek the common organizational principle underlying the novel feature space, by evaluating how the most relevant features clustered together (figure 3).

**Figure 3.**
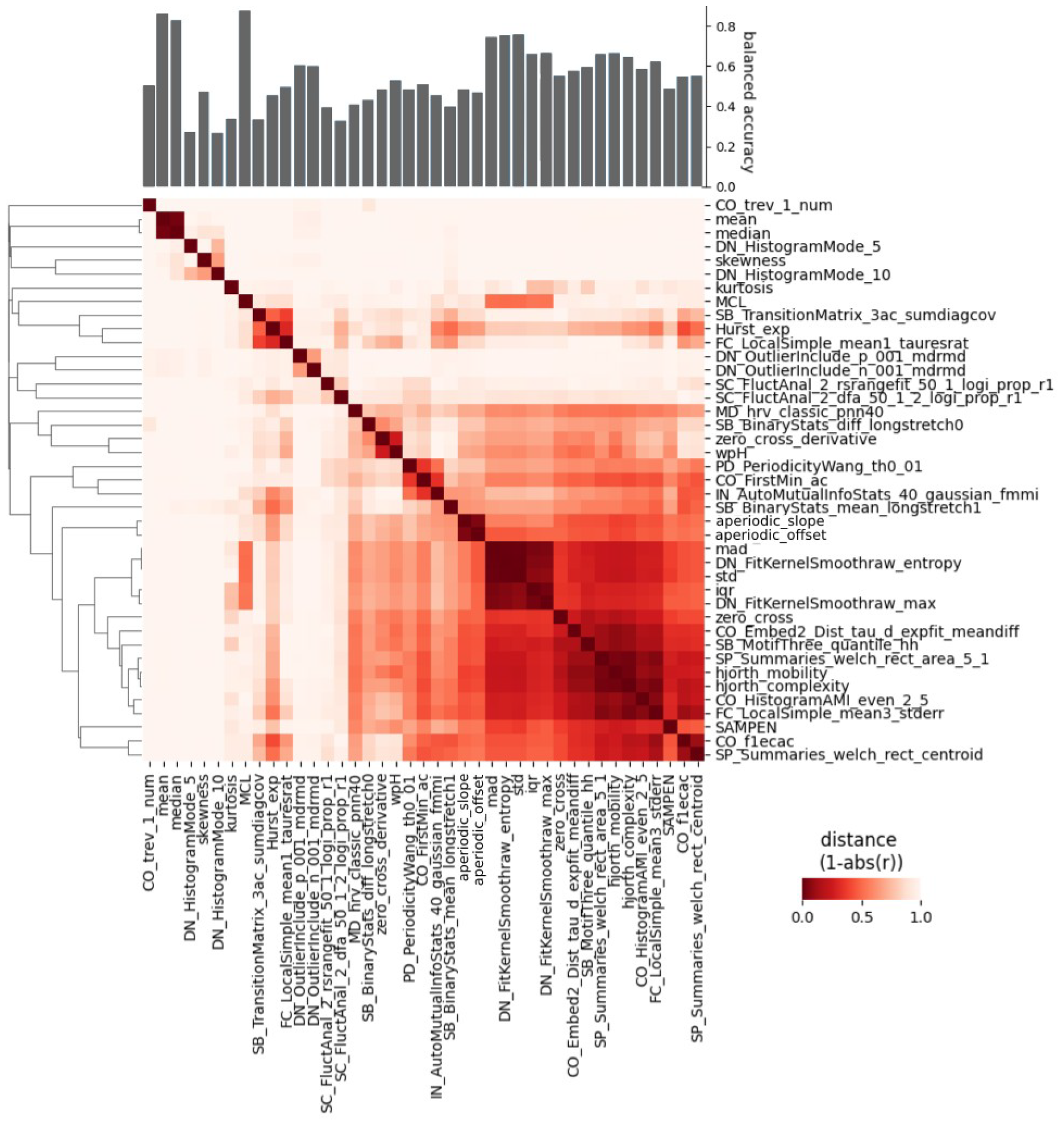
Distance between features computed on broadband signal, represented as correlation matrix in the center. On the left, the dendrogram showing the hierarchical clustering of the features based on the distance. On top of the graph the across subjects classification accuracy for the four classes EC/EO/BSL/VS.

By combining the feature informativeness and the clusters formed based on the distance between pairs of features, we highlight three central subsets of features. The first is the one that pools together mad, std, DN_FitKernelSmoothraw_entropy, and to a slightly lesser extent interquartile range (iqr) and DN_FitKernelSmoothraw_max. All of these features describe the shape and width of the statistical distribution of data.

A second cluster is the one including median and mean, and describing signal’s baseline shifts. This is likely due to slow drifts in the signal which would make inter-block differences between EC and EO relevant.

Intriguingly, the best performing feature in the set (MCL) is only weakly correlated with other features.

### Replication study results

To corroborate the importance of the novel features gathered in the main study, we run a pre-registered replication of the main results on an independent EEG dataset with 210 participants, for the classification of EC/EO (see Methods). Results are in figure 4.

**Figure 4.**
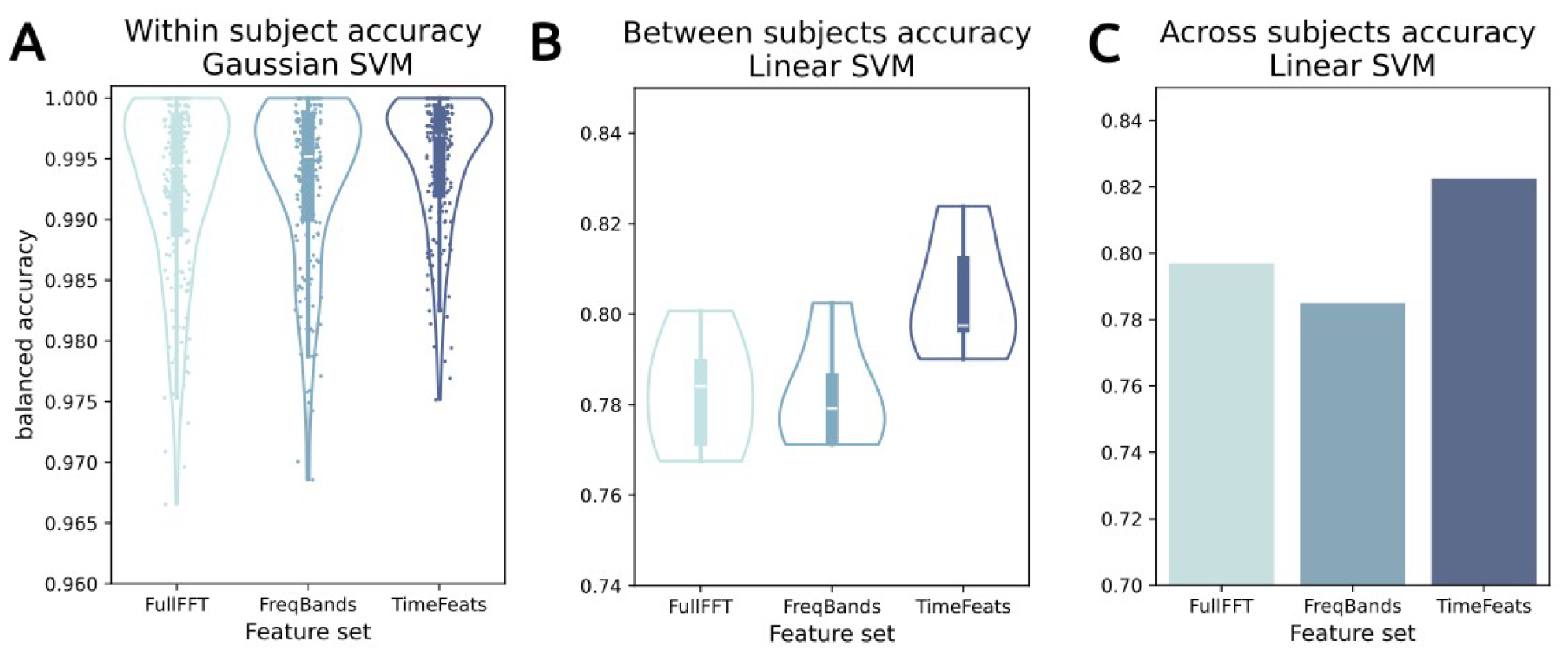
**A**: Within subject classification accuracy, pooled for all subjects. Every point corresponds to a subject. **B**: Between subjects classification accuracy. All subjects left out in each one of the 5 cross-validation folds are pooled together. **C**: Across subjects classification accuracy. Since all subjects are pooled in one train/test set, the analysis yields only one accuracy estimate per Feature set.

Although even in this dataset we observed a marked ceiling effect in accuracy for the “within” subjects classification, we report a significant effect of feature set (F_(2, 418)_=42.959, p<.001, η^2^=.02), with TimeFeats performing best, by a close yet significant margin (TimeFeats=.995; FreqBands & FullFFT=.993; TimeFeats vs FreqBands: t_(209)_=-6.982, p<.001, d=.27). These results were replicated, even more substantially, in the between subjects condition, where the lack of ceiling effect allows a better quantification of the accuracy gain (TimeFeats=.804; FreqBands=.782; FullFFT=.783; TimeFeats vs FullFFT: t_(4)_=-6.48, p=.003, d=1.597), and in the across subjects condition as well (TimeFeats=.822; FreqBands=.797; FullFFT=.784). So all in all, we were able to confirm the relevance of our novel set of features in two independent datasets, from different recording modalities.

## Discussion

In the present work, we characterized states of cortical activity, such as the presence or absence of visual stimulation, by means of a novel multidimensional feature space. By computing the states’ classification accuracy, we found that some of these features, largely neglected in canonical analyses of brain time series, are highly informative for setting brain states apart. Critically, their contribution often exceeds the benchmark defined by classic features such as power changes in canonical frequency bands of interest. Thus, our study paves the way for brain states characterization beyond spectral parameterization.

The most informative feature set broadly describes the variance of the signal, and includes features like standard deviation (std), mean absolute deviation (mad) and interquartile range (iqr). Although these features are related to spectral power it is noteworthy that they are more informative than power. This is especially true in some frequency bands: in the high gamma band, for instance, std yields a better classification accuracy than power. This could be a result of the different algorithms for computing power vs std in frequency bands, with the former relying on FFT and the second band passing the signal to then compute the signal variance: the FFT estimate of power in frequency bands necessarily contains information (such as aperiodic components of the spectrum), perhaps not informative for the current classification.

In this regard, it is of particular interest that in our dataset aperiodic components such as 1/f exponent or offset do not seem particularly informative. This seems in contrast with the established relationship between 1/f exponent and different excitability cortical regimes (33) which should have differentiated between the different brain states examined.

One likely explanation for the poor informativeness of aperiodic components, as much as for slower rhythmic components, is the short duration (1 sec) of the time windows considered here. Previous works, focusing on resting state data, showed that stable power estimates require segments between 30 and 120 seconds (34), and such an estimate was confirmed also for several of the novel features computed here (35). This makes the high classification accuracies obtained in the present study even more remarkable, especially when novel features are considered.

Some of these features, such as Mean Curve Length (MCL), might have been particularly informative compared to others because of the short time window considered. In fact, MCL is computed by taking the average absolute value of the first derivative of the signal. It therefore quantifies the overall absolute change or variability in the signal, irrespective of the sign of change (increase or decrease). A second interesting interpretation of MCL stems from the fact that the process of first order differentiation removes a large part of the 1/f component in the signal (36) so that MCL can be seen as a time-domain analogon of true oscillatory overall power. MCL has also recently been associated with complexity metrics as fractal dimension (37), although in the current work it outperformed several measures of entropy and complexity.

One might still argue that these new features do not add valuable information due to their lack of interpretability. In fact, while we capitalize on decades of research of biophysical models capable of, at least in part, explaining the mechanisms underlying brain oscillations (4,14,38), the study of non-periodic features of electromagnetic brain activity is still in its infancy. Nonetheless, it is rapidly growing, and constantly providing novel insights into the biophysical meaning of these features. A recent work (39) complemented the association between 1/f slope and cortical excitability (33) by showing an inverse, monotonic relationship between 1/f and a measure of signal complexity. Likewise, the importance of variability in the neural signal in determining and predicting behavior is getting increasingly acknowledged (32, for a review), with multiple links between metrics like variability or complexity and arousal being disclosed (31). Furthermore, by using a similar, multidimensional feature approach, Shafiei and colleagues (35) recently described a link between the variation of neurophysiological time dynamics and multiple micro-architectural, structural features.

Moreover, these novel features can increase our understanding of brain oscillations as well. In fact, it is plausible that the list of best performing features could change according to the length of the time window on which the features are computed, leading to crucial insights on the characteristics of brain oscillations at multiple timescales.

The examination of brain activity at different timescales is also a central aspect of brain states (12), since the behavioral readout of brain states exist at different timescales, from milliseconds to days. In this perspective, a richer characterization of brain time series acquires even more importance.

By providing insight on brain states, the novel features discussed here can find applications in a variety of contexts. A successful example is the algorithm for sleep stages classification by (40), which yielded improved accuracy by merging estimates of power in different frequency bands with several features included in the present work, from standard descriptive statistics to nonlinear features such as fractal dimension, permutation entropy and Hjorth parameters of mobility or complexity. This means that novel features can find their relevance directly in the clinical practice.

In conclusion, we advocate for the exploration of novel ways to describe electromagnetic brain activity. Some of these new features are more informative for distinguishing between different brain states than canonical ones based on oscillatory power in predefined frequency bands. These results suggest that a multivariate, data-driven and assumption-free approach to brain time series analysis is a powerful way to study brain states and their transitions.

## Methods

### Study structure

This study consists of a main experiment and a replication experiment. In the main experiment, we analyzed the MEG data of 29 participants (29 males, mean age = 24.9, SD=5.3). These participants performed a battery of tasks, including 5 minutes resting state with eyes closed, (EC), 5 minutes resting state with eyes open (EO) and a speeded detection task introduced by (27) (see *Main experiment data collection*).

Further we aimed at a pre-registered replication experiment of the EC/EO discrimination. For this purpose we relied on an existing open EEG dataset (41), in which 210 participants alternated short sessions of EC and EO resting state. Since the data is publicly available, data handling and preprocessing was performed by a member of the Painlab Munich group. The main authors that performed the pre-registered analysis (Münster’s research group) did not have access to the preprocessed data until the deposit of the pre-registration ([link to osf](https://osf.io/r3976)). Therefore, the authors responsible for the analysis did not have any contact with the data, aside from the information publicly available.

### Main experiment data collection

All 29 subjects provided informed consent in written form prior to their participation in the study. The study protocol was approved by the ethics committee of the Medical Faculty of the University of Münster and the study was conducted according to the Declaration of Helsinki. These participants were recruited from a pool of subjects that had previously participated in MRI experiments at the University of Münster. They hence also provided consent for using the MRI scans in order to allow the source reconstruction in MEG.

Brain magnetic fields were recorded in a magnetically shielded room via a 275 channel whole-head system (OMEGA, CTF Systems Inc, Port Coquitlam, Canada). Data were continuously acquired using a sampling rate of 1200 Hz. Subjects were seated upright, and their head position was comfortably stabilized inside the MEG dewar using pads.

The visual stimuli were projected onto the back of a semi-transparent screen positioned approximately 90 cm in front of the subjects’ nasion using an Optoma EP783S DLP projector with a refresh rate of 60 Hz. The viewing angle ranged from -1.15 to 1.15 in the vertical direction and from -3.86 to 3.86 in the horizontal direction. The subjects’ alertness and compliance were verified by video monitoring.

Participants underwent 5 minutes of resting state with eyes closed (EC), 5 minutes of resting state with eyes open (EO), and a speeded visual detection task. The task was the same as used by (42). In brief each trial started with the presentation of a fixation point. After 500 ms, the fixation point contrast was reduced by 40%, which served as a warning. After another 1500 ms, the fixation point was replaced by a foveal, circular sine wave grating (diameter: 5-; spatial frequency: 2 cycles/deg; contrast: 100%). The sine wave grating contracted toward the fixation point (velocity: 1.6 deg/s) and this contraction accelerated (velocity step to 2.2 deg/s) at an unpredictable moment between 50 and 3000 ms after stimulus onset. The subjects’ task was to press a button with their right index finger within 500 ms of this acceleration. In case the button press occurred within 500 ms after the acceleration point, a positive visual feedback was given to the participants during a resting period of 1000 ms, a negative feedback was given otherwise. The stimulus was turned off after a response was given, or in catch trials 3000 ms after stimulus onset. The paradigm consisted of two blocks of 300 trials in total. At the end of each block, feedback with the percentage of correct responses was provided. The total duration of the paradigm was 7 min.

### MEG Preprocessing and source reconstruction

MEG data were processed using the FieldTrip toolbox (Oostenveld et al., 2011) for MATLAB 2021a (The MathWorks, Inc.) and in-house MATLAB routines. Continuous data were filtered offline with a fourth-order forward-reverse zero-phase Butterworth high-pass filter with a cutoff-frequency of 0.5 Hz and downsampled to 256 Hz for computational efficiency. Independent components (mean number of rejected components per block M = 5; SD = 5.3) arising from heartbeats and eye movements were visually isolated and removed using the runica ICA algorithm.

We used individual T1-weighted MRIs to extract participants’ head model information. First, MRI images were coregisted with the MEG coordinate system via digitized markers in participants’ earnolds using the fieldtrip toolbox (43). Subsequently, subjects’ cortex was described on the surface by means of a cortical sheet through the Freesurfer analysis pipeline (*recon_all* command). Then, cortical sheets were further preprocessed with the Connectome Workbench wb_command (v1.1.1). This resulted in 32492 vertices (neural sources) per hemisphere. To further compute the forward projection matrices (leadfields), we generated a single-shell spherical volume conduction model (44). After computing the leadfields, source activity was estimated on the basis of linear constrained minimum variance (LCMV) beamformer coefficients for each vertex of the cortical sheet (45). For this, sensor covariance was computed across all trials and lambda regularization parameter was set to 5 %. We parcellated the cortical surface according to the HCP-MMP1 atlas [180 parcels per hemisphere; (46)] and from these we chose a subset of 52 parcels marked as visual in the parcellation (see table 2). Within each selected parcel vertex time-series were concatenated and we extracted the first principal component.

In order to prepare the data for classification, we defined 1 s data segments in each condition (EC, EO, BSL and VS). For the EC/EO conditions, we simply segmented the 5 minutes of resting state in each of the conditions. The BSL segments consisted of the 1 s prestimulus window immediately preceding the visual stimulation in each trial, whereas we selected for the VS the time window from 1 to 2 seconds post-stimulus.

### EEG preprocessing

The EEG data was preprocessed using DISCOVER-EEG, an automated preprocessing and feature extraction pipeline to favor the objectivity of the posterior results (47). Briefly, the publicly available dataset consisted of data from 215 subjects (41). Two resting state conditions were acquired in a single session for each participant: eyes open resting state and eyes closed resting state. 8 minutes of each condition were acquired in blocks of 1-minute, interleaving eyes open and eyes closed resting states. Out of the 215 recordings, 1 participant had a corrupted marker file, 1 participant had an empty marker file making it impossible to separate eyes open and eyes closed conditions, 2 participants had truncated data, and 1 participant had a marker file which does not code for the eyes closed condition as disclosed in the original publication. In total, 210 recordings of the eyes closed condition and 211 recordings of the eyes open condition were available for preprocessing. Before preprocessing, the 8 blocks of each condition were extracted and concatenated. Preprocessing was performed separately for eyes open and eyes closed files, keeping the same parameters for both conditions. The one subject missing the eyes closed condition was not included in the further analyses.

### Features selection and sets

All features were computed in MATLAB 2022b (the mathworks). The novel set of features named “TimeFeats” contained a variety of viable descriptors of time series. We tried to keep the list as comprehensive as possible, while capping the features number to ease computability. A detailed description of all the features used can be found at table 1. These included the first 4 moments of probability distributions, measures of entropy (31,32), a set of wide purpose features called *catch22 (28)* tailored to account approximately 90% of the variance explained by the massive amount of features of its mother set *hctsa (29)*, and features previously used in M-EEG, such as Hurst exponent, Hjort parameters and zero crossings (30). This set also included aperiodic components of the power spectrum, as 1/f slope and offset. Since the separation between periodic and aperiodic components can be problematic in data not averaged across repetitions, especially when the data segments are short (1 sec) as in our case, we adopted the following procedure. In each parcel, peaks were found on the average spectra from all the repetitions using the FOOOF algorithm (22). The frequency ranges containing peaks, defined by the central peak frequency and bandwidth obtained by the FOOOF algorithm, were erased from the spectra of every repetition (1 sec segments). Aperiodic estimates of 1/f slope and offsets were obtained by fitting regression lines on the log/log spectra, not including the frequencies and the corresponding power values.

**Table 1:**
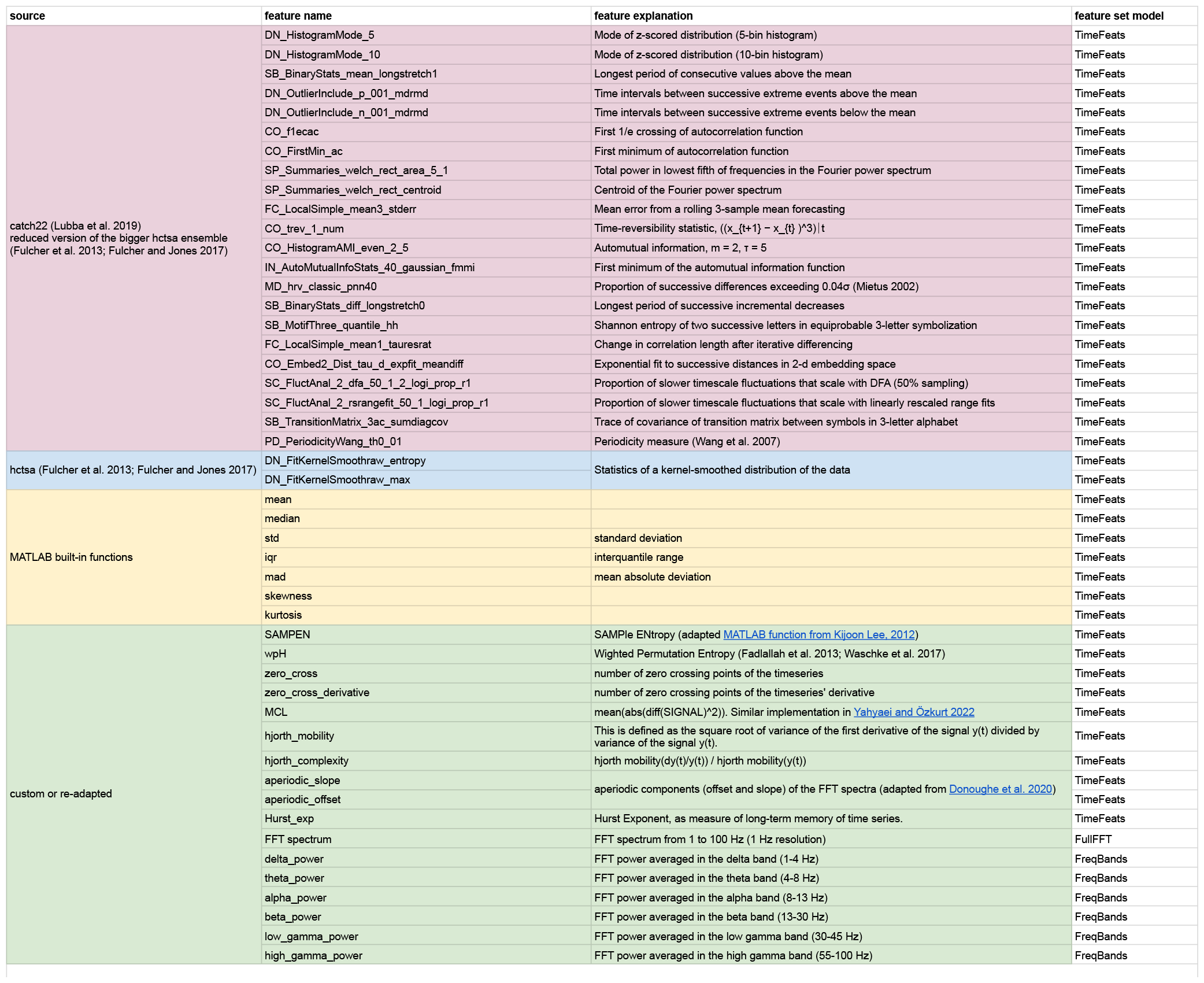
full list of all the features used in the study.

**Table 2:**
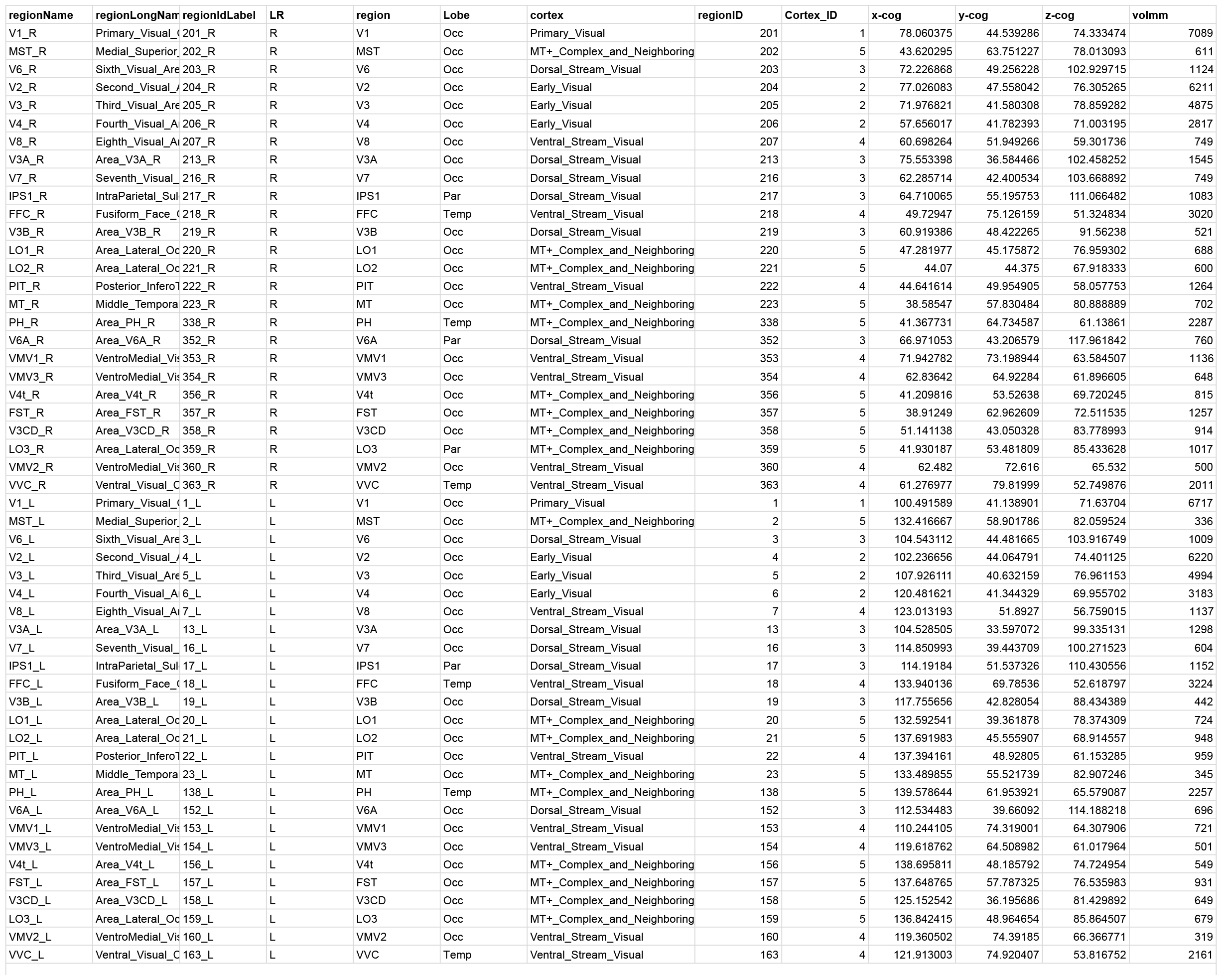
full list of parcels selected in the MEG study.

To have a benchmark of classification accuracy based on spectral power, we created two other sets of features. The first, from hereafter “FreqBands”, contained power estimates averaged in the following frequency bands: delta (1-4 Hz), theta (4-8 Hz), alpha (8-13 Hz), beta (13-30 Hz), low gamma (30-45 Hz) and high gamma (55-100 Hz). The second, named “fullFFT”, contained the full FFT spectrum from 1 to 100 Hz, at a frequency resolution of 1 Hz. Power spectra were computed in fieldtrip, with “mtmfft” method and hanning tapering. The same features were computed on both the MEG and EEG datasets.

For the analyses comparing the classification accuracy of the different feature sets, TimeFeats were computed on the broadband signal (Figure 1, 3 and 4). To get a better overview of the impact of specific features from TimeFeats in different frequency bands, we also computed every feature of TimeFeats in the signal bandpassed in every frequency band (Figure 2).

### Classification and clustering analyses

We compared the overall informativeness of every feature set and of each single feature by computing classification accuracy, either for two-classes (EC-EO, BSL-VS) or all of them combined in a four-classes (EC-EO-BSL-VS) classification task handled with a one-vs-one scheme as default in the Support Vector Classifier from *sklearn*. We adopted different partition schemes to define the training and test sets, defined as follows: “within subject”: one classifier is crossvalidated for each subject; population accuracy is the average of the left-out groups accuracy in the crossvalidation process.

“between subjects”: one classifier is trained on the concatenated data from all subjects minus one, and tested on the left out participant (leave-one-out). This means that none of the subjects in the testing set is represented in the training set. The procedure is repeated iteratively for all subjects, and accuracy is computed on all the subjects left out. In the replication on the independent EEG dataset, given the large number of subjects and in order to cap computation time, we adopted a 5 fold partitioning approach instead of a leave one out (diverging in this way from our pre-registration).

“across subjects”: 85% of the data from each participant will be concatenated across all participants to form the training set, whereas the remaining 15% from each participant will be concatenated to compose the testing set. This means that the subjects in the testing set are also represented in the training set.

Regardless of the partition scheme, the matrix N _observation_ X M _features_ underwent a standard preprocessing in python *sklearn* to standardize values and facilitate the classifier’s convergence. Specifically, missing values along each column were first substituted with the column mean. Then the values of each column were centered around the median and rescaled according to interquantile range (RobustScaler in sklearn). Data was further squeezed to a [-1 1] interval by applying the hyperbolic tangent function to limit even more the effect of outliers. Finally a PCA was applied to select the components accounting for 90% of the total variance before submitting them to a SVC. Default gaussian kernel and C=10 was chosen for all the analyses, except when stated differently: in some cases in fact a linear kernel was chosen to reduce computation time. Classification accuracy was always computed as balanced accuracy (balanced_accuracy_score in sklearn) to account for potential label disbalance in the sample.

In order to evaluate how the single features clustered together, we computed the distance between pairs of features as 1-abs(r), where r is the Pearson correlation coefficient. Hierarchical clustering was performed with complete linkage in scipy. This analysis was performed on the data from the 52 parcels concatenated across subjects.

### Statistical analyses

All inferential statistical analyses, including repeated measures ANOVA and t-test were performed in python with pingouin (v 0.5.3).

## Pre-registration and data availability

The pre-registration for the replication experiment can be found at [OSF](https://osf.io/r3976). Code and data will be made publicly available upon acceptance of the manuscript.

## Acknowledgements

This work was supported by the Deutsche Forschungsgemeinschaft (DFG), GR 2024/11-1 (J.G.).

We acknowledge support from the Open Access Publication Fund of the University of Münster.

